# Group composition and its ontogenetic changes in marine pelagic fishes

**DOI:** 10.1101/2024.04.24.591005

**Authors:** Sho Furuichi, Masahiro Suzuki, Yasuhiro Kamimura, Ryuji Yukami

## Abstract

Group living is a widespread phenomenon among animals and an important trait in fishes. Understanding group composition and its dynamics is fundamental for elucidating ecological and evolutionary processes in natural populations of gregarious animal species, including fishes, but our knowledge of group composition in marine fishes is limited. Here, we examined group composition and its ontogenetic change in two marine fishes, Japanese sardine (*Sardinops melanostictus*) and chub mackerel (*Scomber japonicus*). We found that, at the juvenile stage, groups were composed of individuals with similar body length, age, and growth history. At the sub-adult stage, although body length within a group remained similar, the within-group similarity in age and growth history decreased, approaching the level expected when groups were randomly composed. Therefore, during the early life history, individuals form groups with those spawned at approximately the same time and place (i.e., those from the same batch). As they develop, however, individuals of various origins intermingle and selectively form groups with others exhibiting similar phenotypes. We further found that environmental and geographical factors were also related to the body size and age of individuals distributed. This indicates that Japanese sardine and chub mackerel exhibit phenotype-specific and growth-stage-specific habitat use, which also contributes to the formation and maintenance of structured groups. Our findings provide valuable information on group composition, its ontogenetic change, and its underlying mechanisms in marine fishes, for which the information is limited. These insights may help to clarify ecological and evolutionary processes such as population dynamics, density-dependent phenomena, and the evolution of grouping behaviour in small pelagic fishes.

## 1. INTRODUCTION

Grouping behaviour is widespread across the animal kingdom and is a particularly important trait in fishes. Most fishes form groups throughout—or at least for part of—their life history. There has been an increasing recognition that understanding the social fine structure (i.e., group composition and dynamics) of animal populations is fundamental for understanding ecological and evolutionary processes in natural populations of gregarious animal species, including fishes (Farine et al., 2015).

Group formation provides fishes with many benefits. A key benefit of grouping is a reduced predation risk although there are other benefits, such as facilitating encounters with reproductive partners, improving foraging efficiency, and hydrodynamic gains (Krause & Ruxton, 2002; Pitcher & Parrish, 1993; Ward & Webster, 2016).

Whilst individuals can gain benefits by simply associating with others, the benefits of grouping can be increased by associating with individuals of a particular phenotype (Hoare et al., 2000a; Ranta et al., 1994). For example, the anti-predator benefits of grouping can be enhanced by associating with individuals of the same phenotype (e.g., colour, size, shape, and sex) to reduce a predator’s capture success rate due to the confusion effect (Krakauer, 1995; Landeau & Terborgh, 1986) and to reduce predation risk via the oddity effect (Landeau & Terborgh, 1986; Ohguchi, 1978; Theodorakis, 1989). In addition, grouping benefits can be enhanced by associating with kin and/or familiar individuals (Hamilton, 1964; Ward & Hart, 2003), for example, by increasing reciprocity and behavioural synchronization within a group (e.g., Davis et al., 2017; Hesse et al., 2015), leading to enhanced predator avoidance and evasion (e.g., Chivers et al., 1995; Nadler et al., 2021; Wisenden & Smith, 1998; Wolcott et al., 2017).

Previous investigations have found the composition of free-ranging fish groups to be non-random (Killen et al., 2017; Krause & Ruxton, 2002). In various fish species, groups are assorted by phenotype (e.g., Hoare et al., 2000b; Kelley & Evans, 2018; Krause et al., 1996), and kinship/familiarity assortment has also been reported in some species (e.g., Barber & Ruxton, 2000; Gerlach et al., 2001). However, most of these findings come from studies of freshwater fishes, and information on the group composition of marine fishes is scarce (Hoare et al., 2000a; Krause & Ruxton, 2002), although some clues have been gathered (Sakakura & Tsukamoto, 1997). Generally, habitats of marine fishes are spatially larger than those of freshwater fishes, and groups of marine fishes are more widely and sparsely distributed in the environment, resulting in lower rates of encounters between groups (Croft et al., 2003b). Group encounters are important opportunities for individuals to exchange between groups, and group encounter rates will affect group composition (Croft et al., 2003a, 2003b). Therefore, it is unclear whether the same pattern of group composition occurs in marine fishes as in freshwater fishes.

In addition, traits relevant to group formation, such as mobility, vulnerability to predation, and sociality generally change with growth (Fuiman & Magurran, 1994; Krause & Ruxton, 2002; Ward & Webster, 2016). Therefore, it is possible that group composition changes in accordance with changes in growth stages. However, previous research often examined group composition only at a specific growth stage, thus limiting our understanding of changes in group composition with growth. Group composition and its dynamics are need to be clarified because they are important factors related not only to predation avoidance but also to intra- and inter-specific competition, social structure, information transfer, the spread of pathogens, and thus population and community dynamics (Farine et al., 2015).

The purpose of this study is to clarify group composition and its change with growth in marine fishes. In this study, we focused on Japanese sardine (*Sardinops melanostictus*) and chub mackerel (*Scomber japonicus*), small pelagic fishes that inhabit the western North Pacific around Japan and are commercially and ecologically important fish species because of their high abundance (Yatsu, 2019). Japanese sardine and chub mackerel obligately form groups (shoals), and they sometimes form massive groups consisting of thousands or even a million individuals (Furuichi et al., 2022). Additionally, surveys of these two species are conducted on age-0 fish in different seasons (spring and autumn) by the Japan Fisheries Research and Education Agency, providing data for different growth stages (juvenile and sub-adult).

In this study, we examined the group composition of Japanese sardine and chub mackerel in terms of individual phenotype (body size), age in days, and growth history for juvenile and sub-adult stages. We verified whether the group composition was a random or non-random assortment of individuals. If individuals with similar phenotypes aggregate to form a group, then similarity in body size within the group would be higher than that expected by random aggregation. If individuals spawned at approximately the same time and place—individuals from the same batch (corresponding to kin/familiar)—aggregate to form a group, then similarity in both age and growth history within the group would be higher compared to that expected by random aggregation. Similar growth histories among group members would indicate that they have experienced the same environment over time, meaning a stable group with little turnover of members.

## 2. MATERIALS AND METHODS

### 2.1 Study species

Japanese sardine and chub mackerel inhabit the coastal waters of Japan in the western North Pacific. Both species exhibit similar migration patterns. Adults undertake a northward feeding migration in summer and autumn and a southward spawning/wintering migration in autumn and winter (Kamimura et al., 2021). Spawning mainly occurs off the western and central Pacific coast of Japan from winter to spring (Kanamori et al., 2019; Niino et al., 2021). Most of the planktonic eggs and larvae are transported eastward from the Kuroshio region to the Kuroshio–Oyashio transitional region, where they spend their early growth phase (Kamimura et al., 2015; Niino et al., 2021). Subsequently, juveniles migrate northward to the Oyashio region, which is considered a feeding ground, before returning to the coast of Japan in autumn and winter (Hashimoto et al., 2019; Sakamoto et al., 2019).

### 2.2 Surveys

The Japan Fisheries Research and Education Agency conducts trawl surveys to examine the densities and distributions of small pelagic fishes annually in May and June (spring survey) and September and October (autumn survey) (Figure 1). The spring surveys have been conducted mainly in the Kuroshio–Oyashio transitional region (a nursery area for Japanese sardine and chub mackerel) since 1996, targeting juveniles. The specifications of the trawl net used in the spring surveys are as follows: opening dimensions, 25 m × 25 m; cod-end mesh aperture, 4 mm; tow duration, 30–60 min; ship speed, 3.0–4.0 knots (5.6–7.4 km h^−1^); depth, <25 m. The autumn surveys have been conducted mainly in the Oyashio region (a feeding area for Japanese sardine and chub mackerel) since 2005, targeting sub-adults. The specifications of the trawl net used in the autumn surveys are as follows: opening dimensions, 30 m × 30 m; cod-end mesh aperture, 17.5 mm; tow duration, 15–60 min; ship speed, 3.5–5.0 knots (6.5–9.3 km h^−1^); depth, <40 m. In both surveys, trawl hauls are conducted three times per day. At each station, fish species are identified on board, and the catch is weighed by species. For this study, up to about 400 individuals of each species were randomly selected from the trawl catch and stored at −25 °C.

**FIGURE 1.**
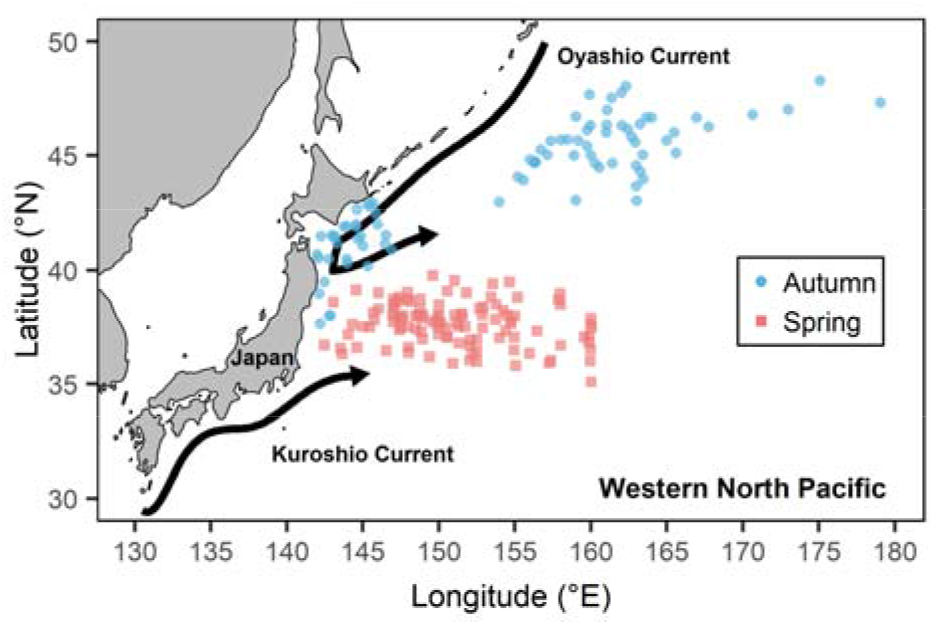
Sampling stations for trawl surveys in the Pacific Ocean off Japan. All sampling stations are shown as red squares for spring (May–June) surveys targeting juveniles and as blue circles for autumn (September–October) surveys targeting sub-adults.

In the laboratory, body size measurement and otolith microstructure analysis were conducted on samples randomly extracted from the frozen specimens to obtain the composition of body size, age in days, and growth history for age-0 fish within each trawl haul. We assumed that a single trawl tow sample was derived from a single shoal or a single cluster (i.e., hereafter, simply ‘group’); a cluster is an aggregate of shoals. Small pelagic fishes are known to form clusters (Haugland & Misund, 2004; Petitgas et al., 2001), and some view clusters as the functional level of fish aggregation (Petitgas, 1996; Reid et al., 2000; Swartzman, 1997).

Up to 50 specimens were randomly selected at each trawl station. Sagittal otoliths were used for otolith microstructure analysis following the methods described by Takahashi et al. (2008) for Japanese sardine and by Takahashi et al. (2014) for chub mackerel. Each Japanese sardine was measured for standard length and each chub mackerel for fork length to the nearest 0.01 mm with digital callipers. Sagittal otoliths of fishes were dissected out and cleaned. The distal side of otoliths was analysed for juvenile and sub-adult Japanese sardine and juvenile chub mackerel, and the dorsal side for sub-adult chub mackerel, as follows. For juvenile and sub-adult Japanese sardine and juvenile chub mackerel, otoliths were mounted on a glass slide with the distal side up using epoxy resin and polished with waterproof sandpaper until the nucleus was clearly visible. For sub-adult chub mackerel, otoliths were initially mounted with the ventral side up on a glass slide using hot water-soluble wax and then polished with waterproof sandpaper until the nucleus was visible. The wax was subsequently removed from the glass slide with hot water, and the otoliths were inverted and remounted on a glass slide with the dorsal side up using epoxy resin. The dorsal side was polished again. Daily growth increments were measured to the nearest 0.1 _μ_m using an otolith measurement system (Ratoc System Engineering, Tokyo, Japan). For both species, the age in days was estimated by adding two to the number of otolith increments because the first increment typically forms 3 days after hatching (Hayashi et al., 1989; Takahashi et al., 2014). In the data analyses below, we used data from years when data of both juvenile and sub-adult stages were available (Japanese sardine: 2014–2017, 2019–2021; chub mackerel: 2006–2013).

### 2.3 Data analysis

To investigate the structure of body length and age composition of groups, we fitted linear mixed models. We estimated the amount of variation associated with among-group differences, as well as differences within groups among individuals, by fitting a random intercept for group ID. The variance of the random intercepts for group ID corresponds to the between-group variance, and the variance of residuals corresponds to the within-group variance. Other factors, such as water temperature and migration, also may contribute to variation in body length and age in days. For example, larger fishes inhabit cooler areas (Lafrance et al., 2005; Morita et al., 2010) and occupy the front of migration (DeBlois & Rose, 1996). Furthermore, yearly fluctuations in growth rate and hatching date have been observed (Furuichi et al., 2020; Niino et al., 2021), which may also contribute to the variation in body length and age in days. Therefore, to compare the within- and between-group variance adjusted for the influence of the spatiotemporal and environmental factors as far as possible (Krause et al., 2000a), the survey year and the sea surface temperature, latitude, and longitude of the survey site were incorporated in the models as fixed effects. In addition, body size and age in days increase over time. To adjust for this effect, the survey date (number of days elapsed since January 1) was also included in the models as a fixed effect.

In the models, survey year and group ID were treated as categorical variables, whereas the others were treated as continuous variables. Response variables and continuous explanatory variables were scaled to zero mean and unit standard deviation to facilitate parameter estimation. In the autumn survey, because there was a strong negative correlation between water temperature and latitude (*r* = −0.86, *p* < 0.001) and they provide the same information, only water temperature was incorporated in the models. The models were fitted using restricted maximum likelihood.

We calculated intraclass correlations (ICCs) to quantify similarity within a group as: ICC = V_G_ / (V_G_ + V_R_), where V_G_ is the among-group variance and V_R_ is the within-group (residual) variance. ICC values necessarily range between 0 and 1. An ICC near 1 indicates high similarity within a group.

To investigate the structure of group composition in terms of growth history, we applied multivariate analysis of variance using distance matrices (Anderson, 2001). In this analysis, response variables were time series of otolith daily growth increments that were square-root-transformed, and their Euclidean distance matrices were calculated. The variance of growth increments in an age tended to increase linearly with increasing mean growth in the age, and therefore square-root transformations were performed on growth increments to homogenize the variances among ages. For the same reasons as for the analyses of body length and age in days, survey year and sea surface temperature, latitude, and longitude of the survey site, as well as group ID, were included in the models as explanatory variables. These models were fitted with type 1 (sequential) sums of squares, which means that each term is sequentially fitted after taking account of the previously fitted terms. This approach allowed us to estimate the effect of group ID after controlling for spatiotemporal and environmental factors. ICCs were also calculated to quantify similarity within a group. We computed the (distance-based) ICCs following the methods of previous studies (Chen & Zhang, 2022; Vogtmann et al., 2017).

To examine whether groups were formed by individuals with similar body size, age in days, and growth history, the observed ICC was compared to the ICCs expected by random grouping (Krause et al., 2000a) in each case for body size, age in days, and growth history. First, a null model was constructed from the above model (alternative model) by excluding group ID and fitted to the data. This model assumes that groups are randomly formed, that is, to be under the null hypothesis. Second, the null distribution of ICCs was obtained by repeating the following procedure 10,000 times: (1) create new data by simulating the response variables (vector or matrix) by bootstrapping the residuals of the fitted null model; (2) fit the alternative model to the new data and calculate ICC. Finally, the observed ICC was compared with the null distribution of ICCs.

All statistical analyses were conducted using R version 4.2.2 (R Development Core Team, 2022). In all analyses, confidence intervals of estimates were evaluated by bootstrapping the residuals (10,000 replicates).

## 3. RESULTS

Data from the juvenile stage (spring surveys) were obtained from 936 individuals in 46 groups of Japanese sardine and from 766 individuals in 65 groups of chub mackerel. Data from the sub-adult stage (autumn surveys) were obtained from 327 individuals in 36 groups of Japanese sardine and from 317 individuals in 48 groups of chub mackerel (Supporting Information Figures S1–S4).

Across all results, Japanese sardine and chub mackerel exhibited similar trends. The within-group similarity (ICC) in body length was higher than that expected when groups were randomly composed, at both the juvenile and sub-adult stages (Figure 2a, b; juveniles and sub-adults of both species, all *p* < 0.001). The within-group similarity in age was higher than that expected when groups were randomly composed at the juvenile stage (Figure 2c, d; both sardine and mackerel, *p* < 0.001), whereas within-group similarity in age substantially decreased from the juvenile to sub-adult stage, approaching the level of similarity expected in randomly composed groups (Figure 2c, d; sardine: *p* = 0.061, mackerel: *p* = 0.004).

**FIGURE 2.**
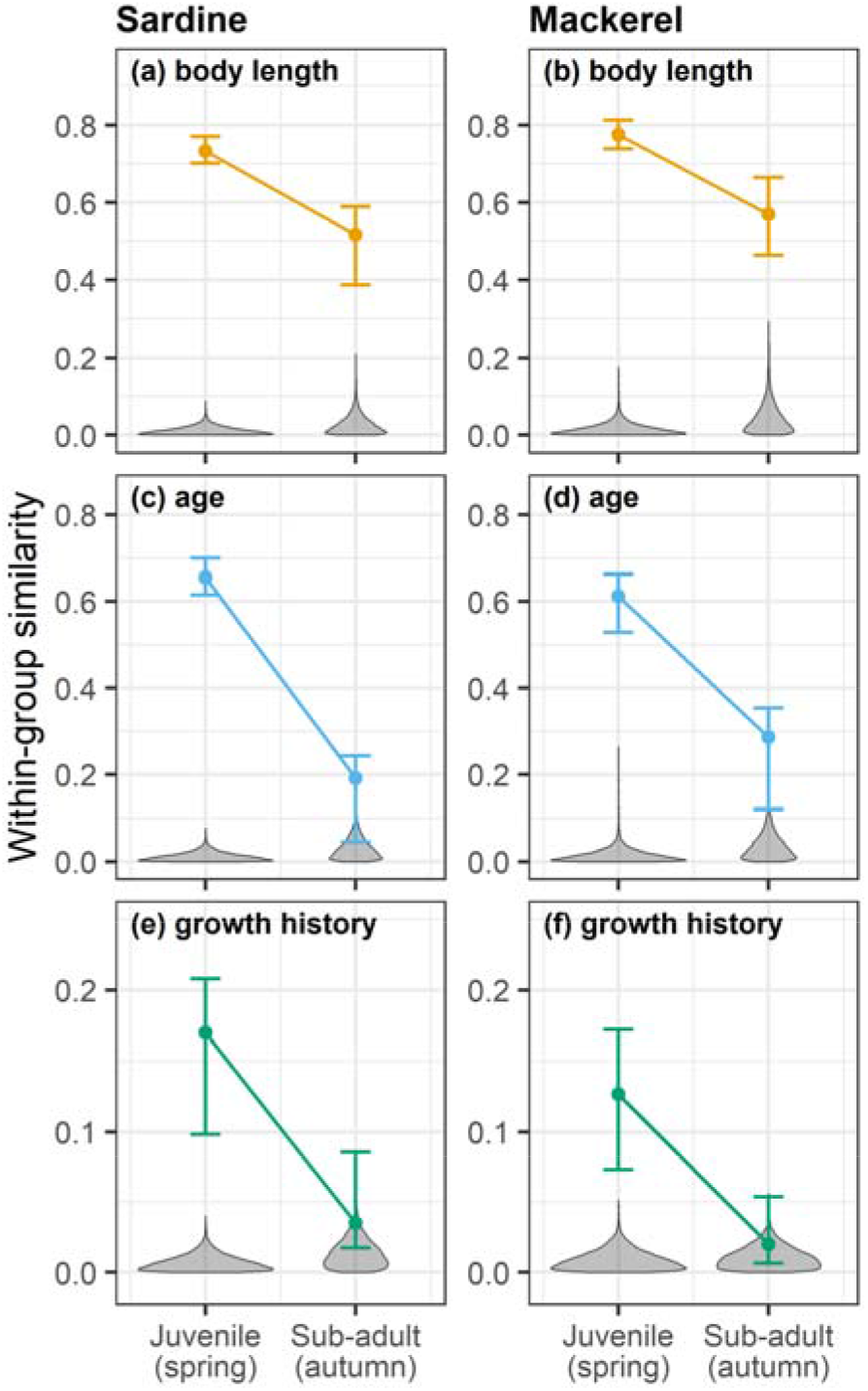
Within-group similarity (intraclass correlation) in (a, b) body length, (c, d) age, and (e, f) growth history at the juvenile and sub-adult stage for Japanese sardine (*Sardinops melanostictus*; left column) and chub mackerel (*Scomber japonicus*; right column). Points represent estimates, and error bars represent 95% confidence intervals. Grey violin plots show the distribution of the similarity expected under a null model of a random assortment of individuals.

The similarity in growth history exhibited patterns like those of similarity in age. At the juvenile stage, the within-group similarity in growth history was higher than that expected when groups were randomly composed (Figure 2e, f; both species, *p* < 0.001). However, at the sub-adult stage, the similarity decreased, reaching levels comparable to those expected when groups were randomly composed (Figure 2e, f; sardine, *p* = 0.57; mackerel, *p* = 0.39).

The fitted models show that environmental variables such as water temperature and spatiotemporal variables such as latitude and longitude were related to body size, age, and growth history of individuals distributed (Figure 3; Table 1). The obvious patterns were that, in both species, larger or older fishes were distributed further east at the juvenile stage, whereas they were distributed further west at the sub-adult stage. Additionally, larger Japanese sardine individuals were distributed further north or in colder environments.

**FIGURE 3.**
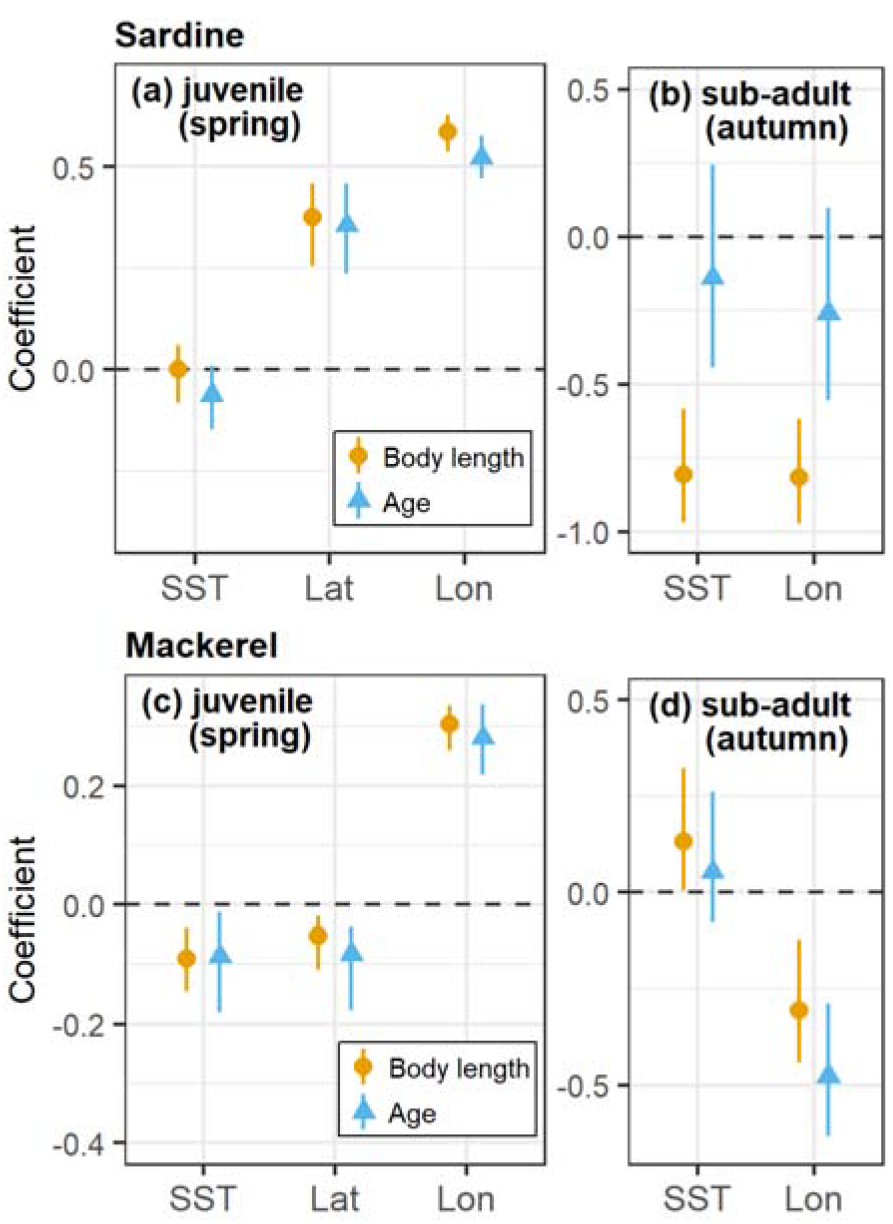
Fixed-effect coefficients from linear mixed models for body length and age at the juvenile and sub-adult stage in Japanese sardine (*Sardinops melanostictus*) and chub mackerel (*Scomber japonicus*). Points represent estimates, and error bars represent 95% confidence intervals. For brevity, only key variables are shown here. Results of all estimates are presented in Supporting Information Tables S1 and S2. SST: sea surface temperature, Lat: latitude, Lon: longitude.

**TABLE 1.** Summary of multivariate analyses of variance using distance matrices for growth history in Japanese sardine (*Sardinops melanostictus*) and chub mackerel (*Scomber japonicus*) at the juvenile and sub-adult stages.

| Growth stage | Source | Sardine |  |  |  | Mackerel |  |  |  |
| --- | --- | --- | --- | --- | --- | --- | --- | --- | --- |
| | | df | SS | MS | $\omega^2$ | df | SS | MS | $\omega^2$ |
| Juvenile | Year | 6 | 195.4 | 32.6 | 0.098 | 7 | 507.5 | 72.5 | 0.142 |
|  | Longitude | 1 | 127.9 | 127.9 | 0.069 | 1 | 11.7 | 11.7 | 0.003 |
|  | Latitude | 1 | 7.0 | 7.0 | 0.003 | 1 | 22.9 | 22.9 | 0.007 |
|  | SST | 1 | 62.8 | 62.8 | 0.035 | 1 | 5.8 | 5.8 | 0.001 |
|  | Station | 36 | 393.9 | 10.9 | 0.163 | 54 | 588.9 | 10.9 | 0.117 |
|  | Residuals | 890 | 1606.5 | 1.8 | — | 701 | 2649.7 | 3.8 | — |
| Sub-adult | Year | 6 | 336.5 | 56.1 | 0.094 | 7 | 821.8 | 117.4 | 0.055 |
|  | Longitude | 1 | 8.4 | 8.4 | 0.000 | 1 | 83.6 | 83.6 | 0.005 |
|  | SST | 1 | 30.7 | 30.7 | 0.008 | 1 | 37.3 | 37.3 | 0.001 |
|  | Station | 27 | 317.6 | 11.8 | 0.031 | 38 | 1399.3 | 36.8 | 0.017 |
|  | Residuals | 291 | 2460.1 | 8.5 | — | 269 | 8657.2 | 32.2 | — |

df: degrees of freedom, SS: type I sums of squares, MS: mean squares, $\omega^2$ : omega-squared
effect sizes, SST: sea surface temperature.

## 4. DISCUSSION

The present study clarifies the group composition and its change with growth in two marine fish species, Japanese sardine and chub mackerel. Understanding the social fine structure (i.e., group composition and dynamics) of animal populations holds fundamental significance for unravelling ecological and evolutionary processes in natural populations of gregarious animal species including fishes (Farine et al., 2015).

Individuals within the same group demonstrated similarities in body length, age, and growth history at the juvenile stage and in body length at the sub-adult stage. Even when accounting for spatiotemporal and environmental factors, the observed within-group similarities surpassed those expected for randomly composed groups, meaning that the high within-group similarities did not merely result from similar environments or circumstances at that time (Krause et al., 2000a). Our results strongly indicate a non-random assortment of individuals among groups of Japanese sardine and chub mackerel.

At the juvenile stage, individuals within a group exhibited similarities in age and growth history as well as body length. This finding indicates that individuals spawned at approximately the same time and place—individuals from the same batch (corresponding to kin/familiar)—form a group. In contrast, at the sub-adult stage, although body lengths within a group remained similar, the similarity in age and growth history decreased an approached the level expected when groups were randomly composed, meaning that individuals of various origins intermingle and selectively form groups with others of similar body length. These findings underscore that the grouping pattern and hence group composition undergo changes corresponding to growth stage.

Despite changes in grouping patterns, phenotypes (body size) remained similar within a group at both the juvenile and sub-adult stages. The processes governing the development of similar phenotypes within a group likely differ between growth stages. At the juvenile stage, individuals spawned at approximately the same time and place formed a stable group; therefore, phenotypic similarity appears to result mainly from passive processes. Phenotypic similarity may arise from continuously sharing the same environment or social conditions, the predation of slow-growing individuals, and the separation of groups due to differences among developmental stages in swimming speeds or ecological preferences along local environmental gradients (Killen et al., 2017). Conversely, at the sub-adult stage, body sizes were similar within a group despite group members originating from various sources, suggesting that phenotypic similarity is primarily due to the active selection for individuals or groups with similar phenotypes. Previous studies have reported the significance of both active and passive processes in generating and maintaining phenotypic similarity within a group (Croft et al., 2003a), which is consistent with the findings of the present study. Furthermore, our findings suggest that the relative importance of each process varies among growth stages.

During their early life history, Japanese sardine and chub mackerel form groups with individuals spawned at approximately the same time and place (kin/familiar). As they develop, individuals of various origins intermingle, and they selectively form groups with individuals exhibiting similar phenotypes. A proximate mechanism that may contribute to this shift in the grouping pattern is the progress in swimming (movement) ability. As individuals develop, their swimming abilities likely improve (Fuiman & Magurran, 1994), leading to increased encounter frequencies with conspecific groups and facilitating the mingling of individuals from diverse origins.

In addition, our results indicate that, at least from the sub-adult stage onwards, phenotypic similarity is a more significant factor in group formation than kinship/familiarity. Previous research has suggested that phenotypic similarity is more important than familiarity in predator avoidance (Cattelan & Griggio, 2020). Given that small pelagic fishes are exposed to high predation pressure from a wide variety of predators (Cury et al., 2000; Overholtz et al., 2007; Szoboszlai et al., 2015), phenotypic similarity is likely to be a more important factor in group formation than kinship/familiarity in Japanese sardine and chub mackerel. In this regard, group size may also be more important than kinship/familiarity (Barber & Wright, 2001). Small pelagic fishes form large groups, which play a crucial role in predator avoidance (Rieucau et al., 2015). Taken together, these results suggest that in Japanese sardine and chub mackerel, as individuals develop, their swimming ability improves and the frequency of encounters with conspecific groups increases, allowing individuals to form large groups with individuals of similar phenotypes.

Such an ontogenetic shift in grouping pattern might be a prevalent phenomenon among small pelagic fishes, whether freshwater or marine, because the improvement of swimming ability with growth is common in fishes (Fuiman & Magurran, 1994), and small pelagic fishes are typically subject to high predation pressure from a diverse array of predators (Cury et al., 2000; Overholtz et al., 2007; Szoboszlai et al., 2015). Indeed, while group formation among related individuals has been reported in the wild, kin grouping is documented primarily during early life history and becomes rare in adults (Ward et al., 2020). In addition, it has been also reported that adult fishes prioritize phenotypic homogeneity over familiarity in group formation in the Mediterranean killifish (Cattelan & Griggio, 2020).

Phenotype-specific and growth-stage-specific habitat use restricts encounters between individuals or groups with different phenotypes, contributing to the maintenance of group structure (Croft et al., 2003a; Krause et al., 2000b). In the present results, environmental (e.g., water temperature) and geographical factors (e.g., latitude and longitude) were found to be related to the body size and age of individuals distributed. For instance, larger or older fishes were distributed further east at the juvenile stage and further west at the sub-adult stage. Additionally, larger sardines were distributed in colder or more northern environments. These results are consistent with the migration patterns of Japanese sardine and chub mackerel (see section 2.1) and the habitat preference suggested in a previous study (Sakamoto et al., 2022). Thus, our results indicate that Japanese sardine and chub mackerel exhibit phenotype-specific and growth-stage-specific habitat use, and these habitat uses likely contribute to the formation and maintenance of structured groups in these species. Overall, our results reveal that the population structure of Japanese sardine and chub mackerel is structured at the coarse scale by phenotype-specific and growth-stage-specific habitat use and at the fine scale by grouping behaviour through active and passive processes.

In this study, we assumed that a single trawl tow sample was derived from one group (single shoal or cluster). In reality, however, it is possible that more than one group was included in a single trawl tow sample. Although the potential impact on our results would be small, the within-group similarity estimated in this study might be underestimated.

Based on our findings, we conclude that group composition is non-random in Japanese sardine and chub mackerel, and group composition changes according to growth stage. During early life history, these fishes formed groups with individuals that had been spawned at approximately the same time and place (kin/familiar). But as they developed, individuals of various origins intermingled, and they came to selectively form groups with others having similar phenotypes. Furthermore, phenotype-specific and growth-stage-specific habitat use also contribute to the formation and maintenance of structured groups in these species. This study provides valuable information on group composition, its dynamics, and its underlying mechanisms in marine fishes, for which the information is limited. This information can help to clarify ecological and evolutionary processes such as population dynamics, density-dependent phenomena, and the evolution of grouping behaviour in small pelagic fishes.

## AUTHOR CONTRIBUTIONS

R.Y. supervised the surveys. S.F. conceived of the study and performed data analysis. S.F., Y.K., M.S., and R.Y. contributed to data collection. S.F. wrote the manuscript with support from Y.K., M.S., and R.Y. All authors discussed the results and contributed to the final manuscript.

## ACKNOWLEDGEMENTS

The study was made possible by enormous efforts undertaken by numerous researchers, as well as the captains, officers, and crews of the training vessel *Hokuho-Maru* and research vessel *Soyo-Maru*. We thank all members of our laboratory for their helpful discussions and assistance with data collection and M. Takahashi of the Japan Fisheries Research and Education Agency for his help with otolith analyses in chub mackerel. We also thank English-speaking professional editors from ELSS, Inc., for English proofreading. The findings and conclusions of this article are the sole responsibility of the authors and do not represent the official views of the Japan Fisheries Research and Education Agency.

