## Supporting Information for "Group composition and its ontogenetic changes in marine pelagic fishes"

Table S1. Estimates and 95% confidence intervals (CIs) for parameters of linear mixed models for body length and age at the juvenile and sub-adult stages in Japanese sardine (*Sardinops melanostictus*).

| Growth stage | Parameter | Body length | Age |
|---|---|---|---|
| | | Estimate (95% CI) | Estimate (95% CI) |
| Juvenile | Intercept | −0.39 (−0.54, −0.28) | −0.51 (−0.66, −0.38) |
| | Longitude | 0.59 (0.54, 0.63) | 0.52 (0.47, 0.58) |
| | Latitude | 0.38 (0.25, 0.46) | 0.36 (0.24, 0.46) |
| | Sea surface temperature | 0.00 (−0.08, 0.06) | −0.06 (−0.15, 0.01) |
| | Catch date | −0.01 (−0.16, 0.21) | −0.01 (−0.20, 0.20) |
| | $Year_{2015}$ | 0.15 (−0.02, 0.28) | 0.41 (0.23, 0.58) |
| | $Year_{2016}$ | 1.11 (1.00, 1.25) | 1.20 (1.06, 1.36) |
| | $Year_{2017}$ | 0.22 (0.04, 0.48) | 0.28 (0.06, 0.54) |
| | $Year_{2019}$ | 0.06 (−0.16, 0.36) | 0.38 (0.10, 0.68) |
| | $Year_{2020}$ | 0.49 (0.26, 0.79) | 0.68 (0.38, 1.00) |
| | $Year_{2021}$ | 0.73 (0.45, 1.15) | 0.79 (0.42, 1.19) |
| | $\sigma_{group}$ | 0.82 (0.77, 0.86) | 0.82 (0.76, 0.86) |
| | $\sigma_{residual}$ | 0.50 (0.45, 0.52) | 0.60 (0.54, 0.62) |
| Sub-adult | Intercept | 0.58 (0.42, 0.75) | 0.35 (0.05, 0.65) |
| | Longitude | −0.82 (−0.97, −0.62) | −0.26 (−0.55, 0.10) |
| | Sea surface temperature | −0.81 (−0.97, −0.58) | −0.14 (−0.44, 0.24) |
| | Catch date | 0.19 (0.11, 0.29) | 0.67 (0.51, 0.82) |
| | $Year_{2015}$ | −0.11 (−0.38, 0.12) | −1.07 (−1.53, −0.62) |
| | $Year_{2016}$ | −0.06 (−0.35, 0.18) | −0.71 (−1.28, −0.29) |
| | $Year_{2017}$ | −1.01 (−1.26, −0.71) | −0.62 (−1.09, −0.08) |
| | $Year_{2019}$ | −1.24 (−1.54, −0.95) | 0.17 (−0.37, 0.66) |
| | $Year_{2020}$ | −0.65 (−0.96, −0.40) | −0.12 (−0.66, 0.39) |
| | $Year_{2021}$ | −1.18 (−1.56, −0.88) | −0.17 (−0.82, 0.37) |
| | $\sigma_{group}$ | 0.67 (0.50, 0.71) | 0.59 (0.26, 0.64) |
| | $\sigma_{residual}$ | 0.65 (0.56, 0.67) | 1.21 (1.05, 1.27) |

Table S2. Estimates and 95% confidence intervals (CIs) for parameters of linear mixed models for body length and age at the juvenile and sub-adult stages in chub mackerel (*Scomber japonicus*).

| Growth stage | Parameter | Body length | Age |
|---|---|---|---|
| | | Estimate (95% CI) | Estimate (95% CI) |
| Juvenile | Intercept | −0.64 (−0.75, −0.52) | −0.41 (−0.58, −0.20) |
| | Longitude | 0.30 (0.26, 0.34) | 0.28 (0.22, 0.34) |
| | Latitude | −0.05 (−0.11, −0.02) | −0.08 (−0.18, −0.04) |
| | Sea surface temperature | −0.09 (−0.15, −0.04) | −0.09 (−0.18, −0.01) |
| | Catch date | 0.52 (0.48, 0.63) | 0.60 (0.53, 0.77) |
| | $\mathrm{Year}_{2007}$ | 0.67 (0.54, 0.83) | 0.46 (0.24, 0.74) |
| | $\mathrm{Year}_{2008}$ | 0.30 (0.17, 0.47) | 0.24 (0.03, 0.52) |
| | $\mathrm{Year}_{2009}$ | 1.30 (1.15, 1.49) | 1.22 (0.97, 1.53) |
| | $\mathrm{Year}_{2010}$ | 0.54 (0.34, 0.68) | 0.40 (0.09, 0.63) |
| | $\mathrm{Year}_{2011}$ | 0.01 (−0.16, 0.17) | 0.17 (−0.08, 0.45) |
| | $\mathrm{Year}_{2012}$ | 0.58 (0.39, 0.69) | −0.12 (−0.40, 0.07) |
| | $\mathrm{Year}_{2013}$ | 1.63 (1.45, 1.74) | 1.26 (0.98, 1.45) |
| | $\sigma_{group}$ | 0.52 (0.47, 0.54) | 0.59 (0.50, 0.62) |
| | $\sigma_{residual}$ | 0.28 (0.25, 0.29) | 0.47 (0.42, 0.49) |
| Sub-adult | Intercept | −0.95 (−1.29, −0.60) | −1.13 (−1.38, −0.66) |
| | Longitude | −0.31 (−0.44, −0.12) | −0.48 (−0.63, −0.29) |
| | Sea surface temperature | 0.13 (0.01, 0.32) | 0.05 (−0.08, 0.26) |
| | Catch date | 0.28 (0.13, 0.38) | 0.32 (0.27, 0.48) |
| | $\mathrm{Year}_{2007}$ | 1.38 (0.90, 1.70) | 1.17 (0.70, 1.52) |
| | $\mathrm{Year}_{2008}$ | 1.15 (0.69, 1.66) | 1.12 (0.42, 1.40) |
| | $\mathrm{Year}_{2009}$ | 1.03 (0.62, 1.48) | 1.32 (0.74, 1.63) |
| | $\mathrm{Year}_{2010}$ | 1.22 (0.80, 1.67) | 1.55 (0.96, 1.86) |
| | $\mathrm{Year}_{2011}$ | 0.49 (0.10, 0.91) | 0.82 (0.32, 1.19) |
| | $\mathrm{Year}_{2012}$ | 1.77 (1.33, 2.14) | 1.73 (1.17, 2.07) |
| | $\mathrm{Year}_{2013}$ | 0.51 (0.14, 0.87) | 1.33 (0.84, 1.63) |
| | $\sigma_{group}$ | 0.68 (0.54, 0.73) | 0.45 (0.25, 0.48) |
| | $\sigma_{residual}$ | 0.59 (0.48, 0.62) | 0.71 (0.61, 0.74) |

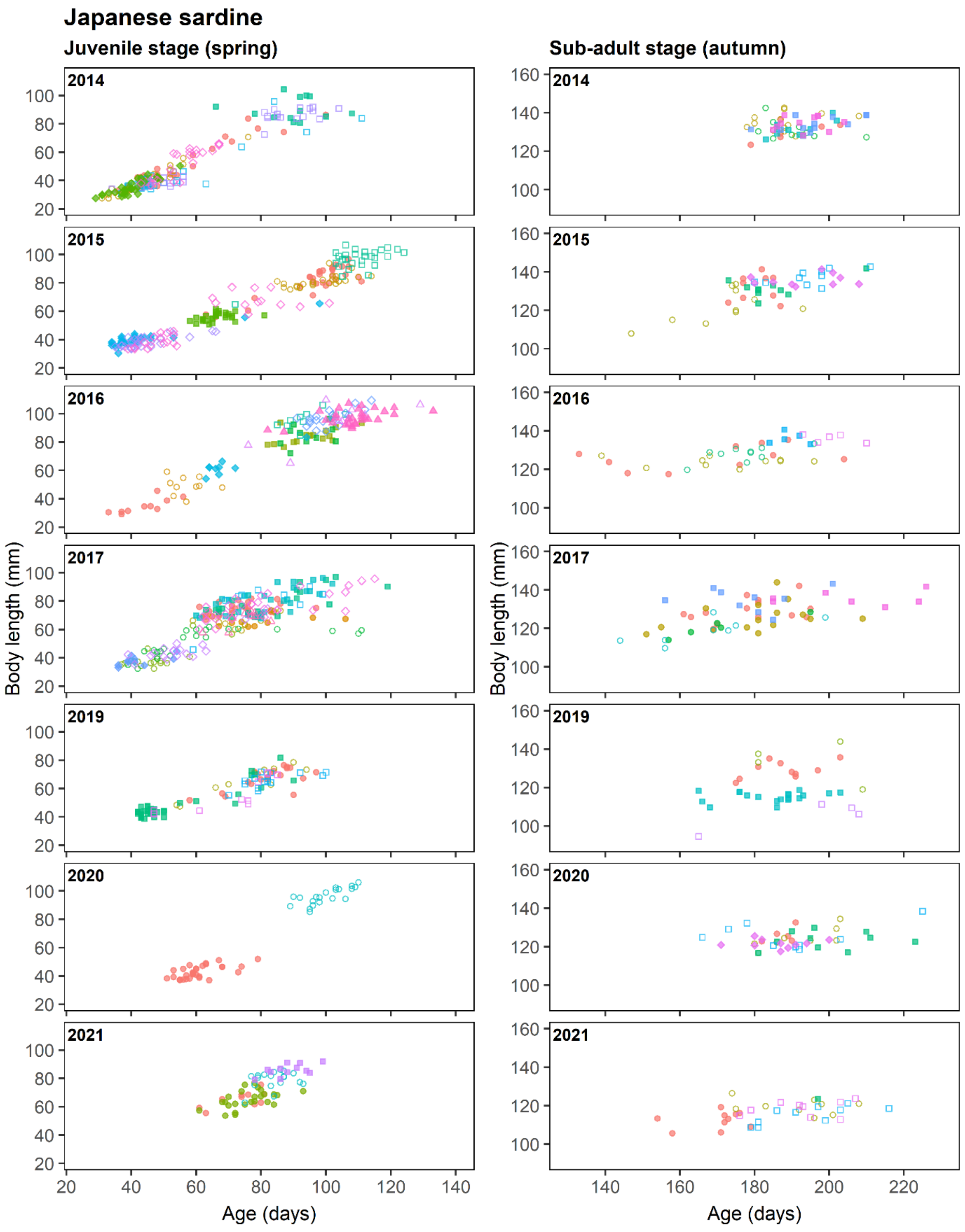


Figure S1. Scatter plots for age and body length at the juvenile stage (left column) and sub-adult stage (right column) in Japanese sardine (*Sardinops melanostictus*) in each year. Each point represents an individual. Different symbols represent different groups.

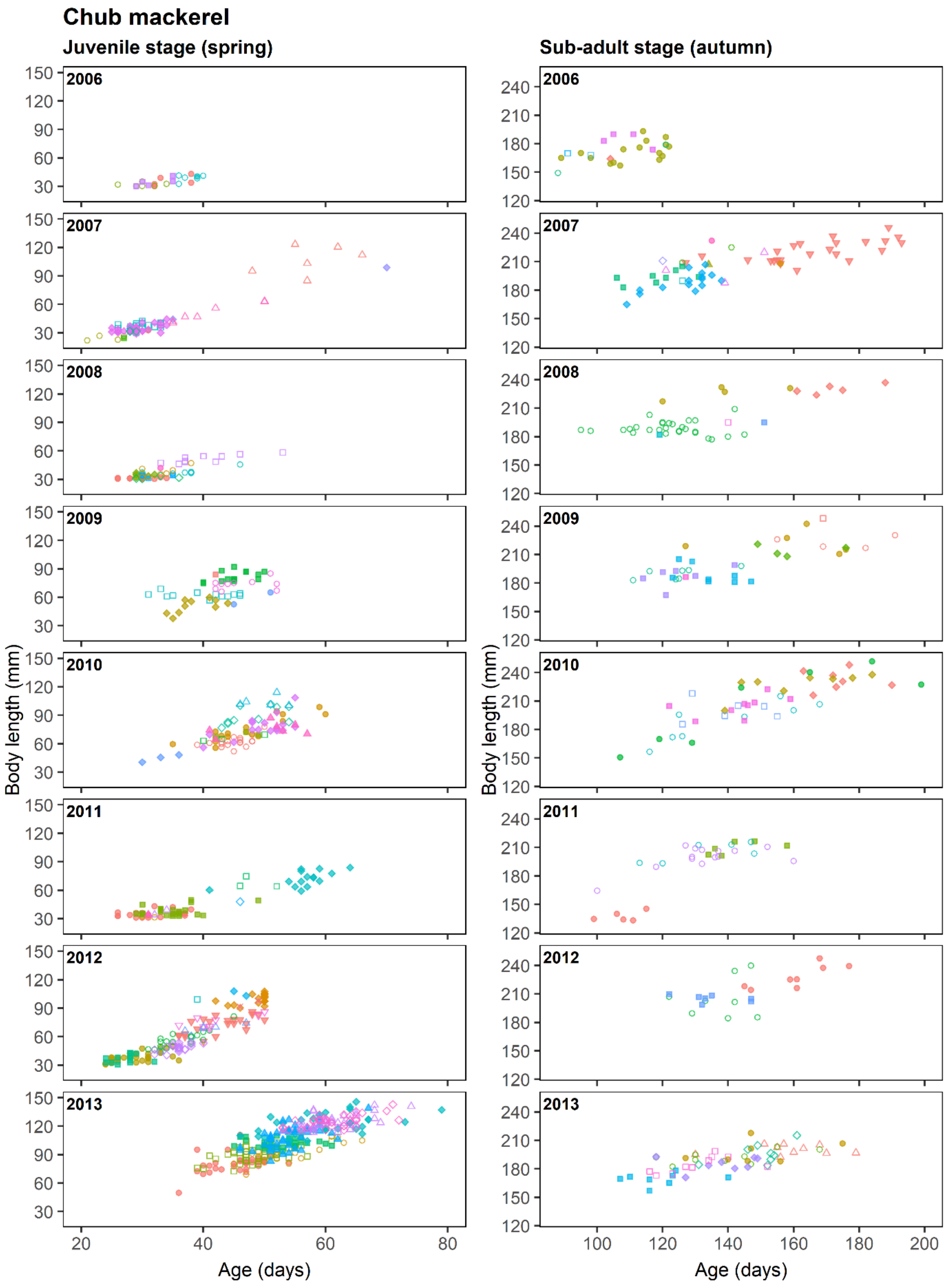


Figure S2. Scatter plots for age and body length at the juvenile stage (left column) and sub-adult stage (right column) in chub mackerel (*Scomber japonicus*) in each year. Each point represents an individual. Different symbols represent different groups.

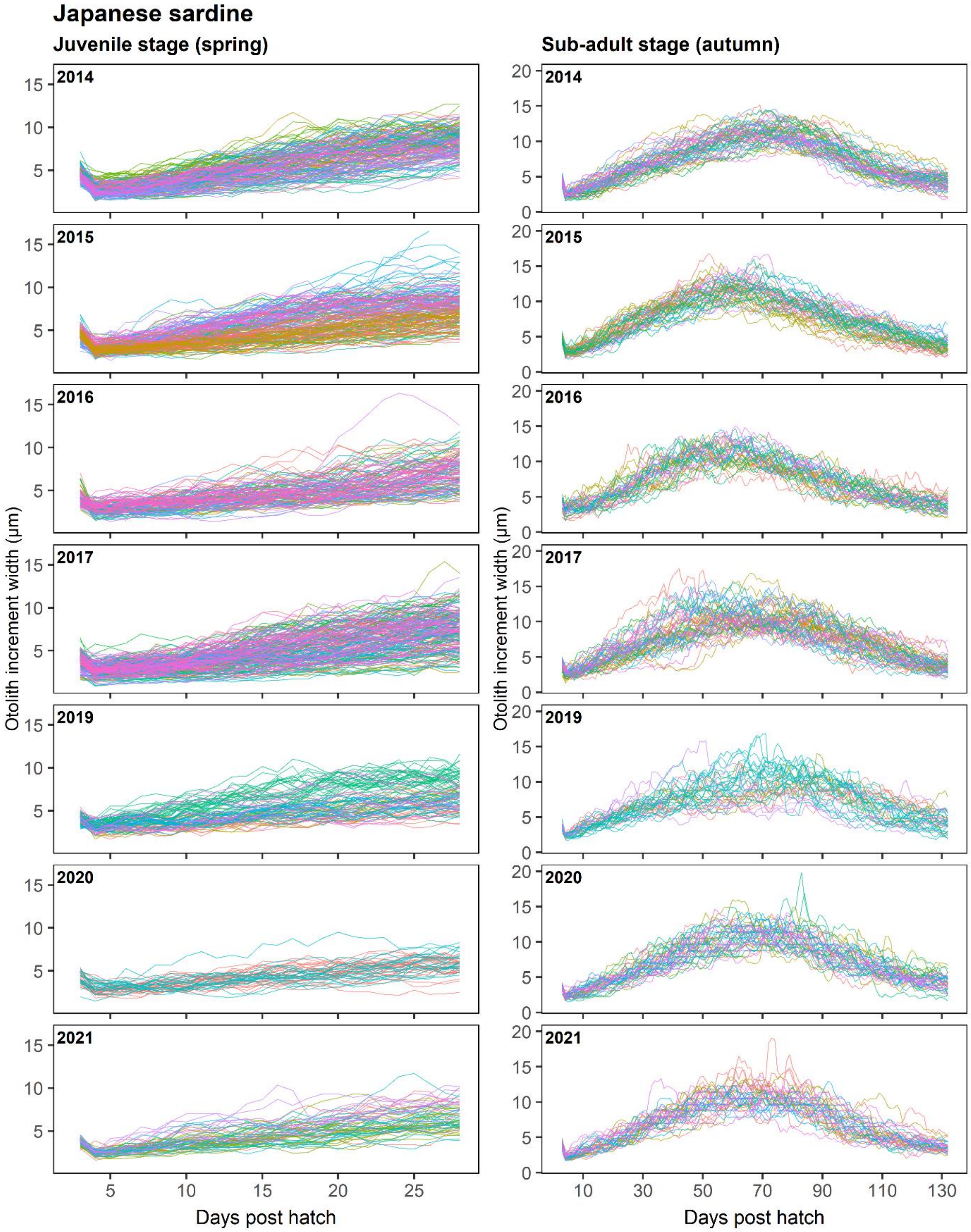


Figure S3. Growth history at the juvenile stage (left column) and sub-adult stage (right column) in Japanese sardine (*Sardinops melanostictus*) in each year. Each line represents an individual. Different colours represent different groups.

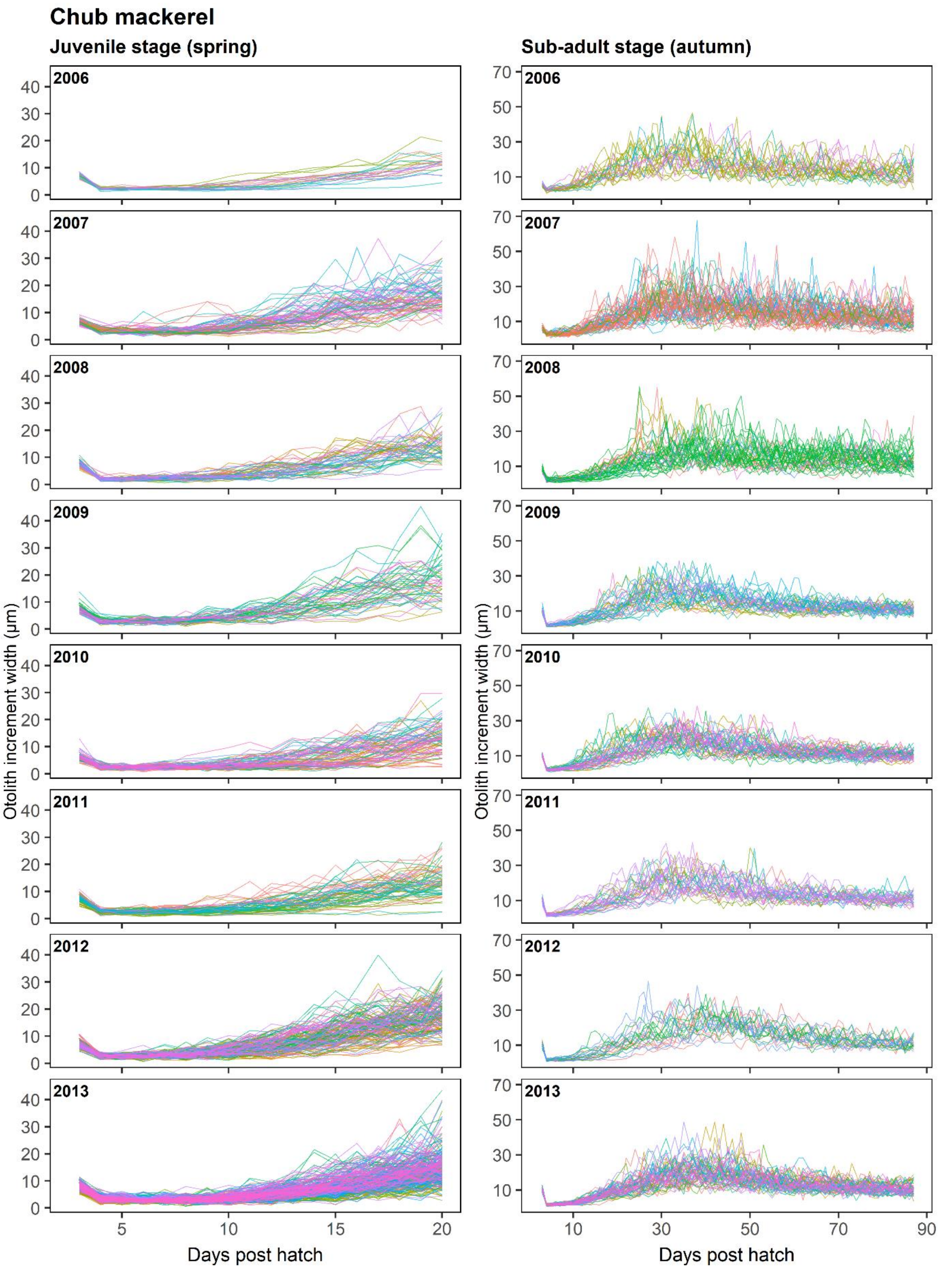


Figure S4. Growth history at the juvenile stage (left column) and sub-adult stage (right column) in chub mackerel (*Scomber japonicus*) in each year. Each line represents an individual. Different colours represent different groups.
